# Combinatorial pathway engineering using multiplex Serine recombinase-Assisted Genome Engineering (mSAGE)

**DOI:** 10.1101/2025.07.11.664204

**Authors:** Jay D. Huenemann, Austin L. Carroll, Joshua R. Elmore, Gara N. Dexter, Thom D. Mand, Dawn M. Klingeman, William G. Alexander, Adam M. Guss

**Author notes:** Notice: This manuscript has been authored by UT-Battelle, LLC, under Contract No. DE-AC0500OR22725 with the U.S. Department of Energy. The United States Government retains and the publisher, by accepting the article for publication, acknowledges that the United States Government retains a non-exclusive, paid-up, irrevocable, world-wide license to publish or reproduce the published form of this manuscript, or allow others to do so, for the United States Government purposes. The Department of Energy will provide public access to these results of federally sponsored research in accordance with the DOE Public Access Plan (http://energy.gov/downloads/doe-public-access-plan).

## Abstract

Low efficiency of DNA integration into the chromosome limits multiplexed bacterial genetic engineering. We address this by establishing multiplexed Serine recombinase-Assisted Genome Engineering (mSAGE), enabling simultaneous, site-specific insertion of three DNAs into the chromosome of *Pseudomonas putida* KT2440. We construct a combinatorial library of isophthalate catabolism and transport genes through a single, pooled transformation of three gene libraries, rapidly identifying the best combination of genes for biodegradation of this plastic comonomer.

---

Genetic engineering of microbes is a promising route to develop biotechnological solutions for societal challenges in energy, resilient agriculture, environmental pollution, human healthcare, and sustainable chemical production [1-4]. Non-model microbes have beneficial properties that make them ideal hosts for utilizing low-cost feedstocks, such as the aromatic molecules found in lignin and plastic wastes, or for producing challenging bioproducts. However, these same microbes lack tools for high-throughput genome engineering [5] especially for targeted insertion of DNA into the chromosome. These tools also have drawbacks, with homologous recombination and recombineering approaches typically having low transformation efficiency [6]. These methods also do not allow multiplexed genome engineering, limiting their applications for high-throughput, combinatorial screening.

Serine recombinase-Assisted Genome Engineering (SAGE) is an efficient, organism-agnostic chromosomal engineering tool that addresses these issues [7]. Transient expression of a sitespecific serine recombinase allows for integration of a cargo vector containing a short *attP* sequence into a genome containing a corresponding *attB* sequence. The original SAGE system allows chromosomal insertion of up to 10 constructs via orthogonal recombinases, and additional recombinases have been discovered in nature with orthogonal *attB* and *attP* sites [6-12], allowing future expansion of SAGE.

We sought to leverage the high efficiency of SAGE and the orthogonality of serine recombinases to enable multiplexed genome engineering, called multiplexed SAGE (mSAGE). In this process, multiple genetic cargos containing distinct *attP* sites are introduced to a single strain at distinct loci harboring the corresponding *attB* site. This can be achieved in a single transformation step, and by introducing libraries of *attP* cargo vectors, mSAGE enables highly efficient combinatorial library generation and screening from a single transformation (Figure 1).

**Figure 1.**
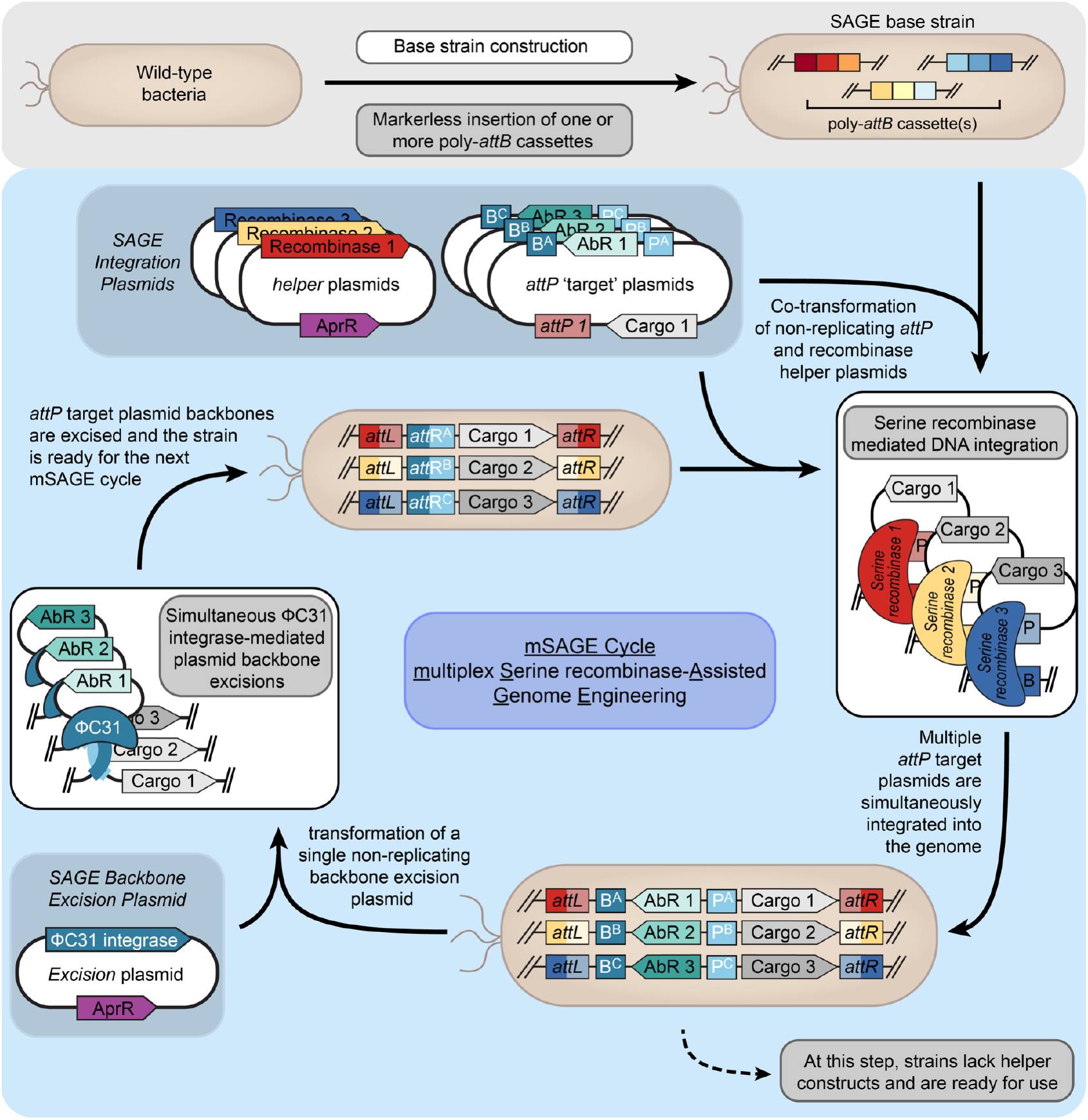
mSAGE approach for multiplexed insertion of DNA into the *P. putida* chromosome. The poly-*attB* harboring strain, AG5577 can be transformed at high efficiency with three recombinases and corresponding cargo DNA pools. Antibiotic selection and outgrowth produce genomically integrated DNA at three distinct sites; the recombinase expressing vector does not replicate in AG5577 and is lost. Antibiotic resistance markers and cargo DNA elements required for maintenance in *E. coli* (but not in *P. putida*) can be removed by transformation with a temperature sensitive vector expressing PhiC31. Heat curing of this plasmid results in a strain ready for additional SAGE using the same antibiotic resistance markers and orthogonal recombinases.

To enable this system, we tested nine orthogonal recombinases in *Pseudomonas putida* by first inserting the *attB* sites for the nine recombinases, evenly split into three landing pads flanked by rho-independent terminators, into the *P. putida* chromosome at three neutral loci: *hsdR, ampC*, and 3’ of the *fvpA* gene, resulting in strain AG5577 (Supplemental Figure S1). We then tested DNA insertion efficiency by co-transforming AG5577 with a non-replicating recombinase-expression plasmid and a non-replicating SAGE integration vector. Six recombinases enabled integration of DNA into the genome with high efficiency (≥10^7^ /µg DNA), with three also having 100% insertional accuracy (insertion into the target *attB* site) (Supplementary Figure S1). Additionally, we demonstrated that another recombinase (PhiC31) expressed from a *P. putida* temperature-sensitive vector (pGW30) could be used to remove antibiotic resistance cassettes introduced by SAGE, allowing for the recycling of these markers in further rounds of SAGE [7]. Because of the exceptionally high efficiency of integration, we tested whether SAGE could be used for multiplexing. A library of SAGE integration vectors was created that combine each of three different antibiotic resistance markers (kanamycin, gentamicin, and streptomycin/spectinomycin) with *attP* sites for each recombinase (Supplemental Figure S3). These cargo vectors also contain rho-independent terminators to insulate the cargo from surrounding genes and PhiC31 *attB* and *attP* sites to allow for resistance marker removal. Co-transformation of these three cargo plasmids and the three corresponding recombinase expression plasmids in a single electroporation cuvette resulted in ≥10^6^ transformants per µg DNA, with all three cargo plasmids simultaneously inserting into the correct locus (Supplemental Figure S4). Further transformation of the PhiC31 recombinase expression plasmid resulted in simultaneous excision of all three antibiotic resistance genes (Supplemental Figure S5), preparing the strain for additional DNA insertions.

As a demonstration of the utility of highly efficient, multiplexed insertion of heterologous DNA, we used mSAGE to find an optimal combination of genes for catabolism of isophthalic acid (IPA, structure in Figure 2B) in *P. putida*. IPA is a non-native carbon source that is significant component of some commercial formulations of polyethylene terephathlate, a major component of plastics waste. A pathway for IPA catabolism in *Comamonas* sp. E6 has been previously described (Supplemental Figure S6) [13]. We developed a library of candidate genes for each step of IPA import and catabolism using genes from *Comamonas* sp. E6 and by isolating IPA-consuming bacteria from the environment and performing *in silico* analyses (BLAST alignment and genomic context) of their genomes to identify orthologs of the *iphAD* (dioxygenase), *iphB* (dehydrogenase) and putative IPA transport genes. We also included candidate orthologs based on BLAST alignments with the published genome sequences for *Burkholderia* sp. CCGE10002 (Genbank accession number CP002013, CP002014, CP002015, and CP002016) [14] and *Maritimibacter alkaliphilus* HTCC2654 (ENA accession number GCA_000152805) [15] (Supplementary Table S3, Supplemental Figure S6). For use in mSAGE, expression cassettes for six orthologs of *iphAD*, six orthologs of *iphB*, and five putative transporters were designed to simultaneously integrate into *P. putida* strain AG5577 at the Bxb1, R4, and TG1 *attB* sites, respectively (Figure 2A, Supplementary Figure S4).

**Figure 2.**
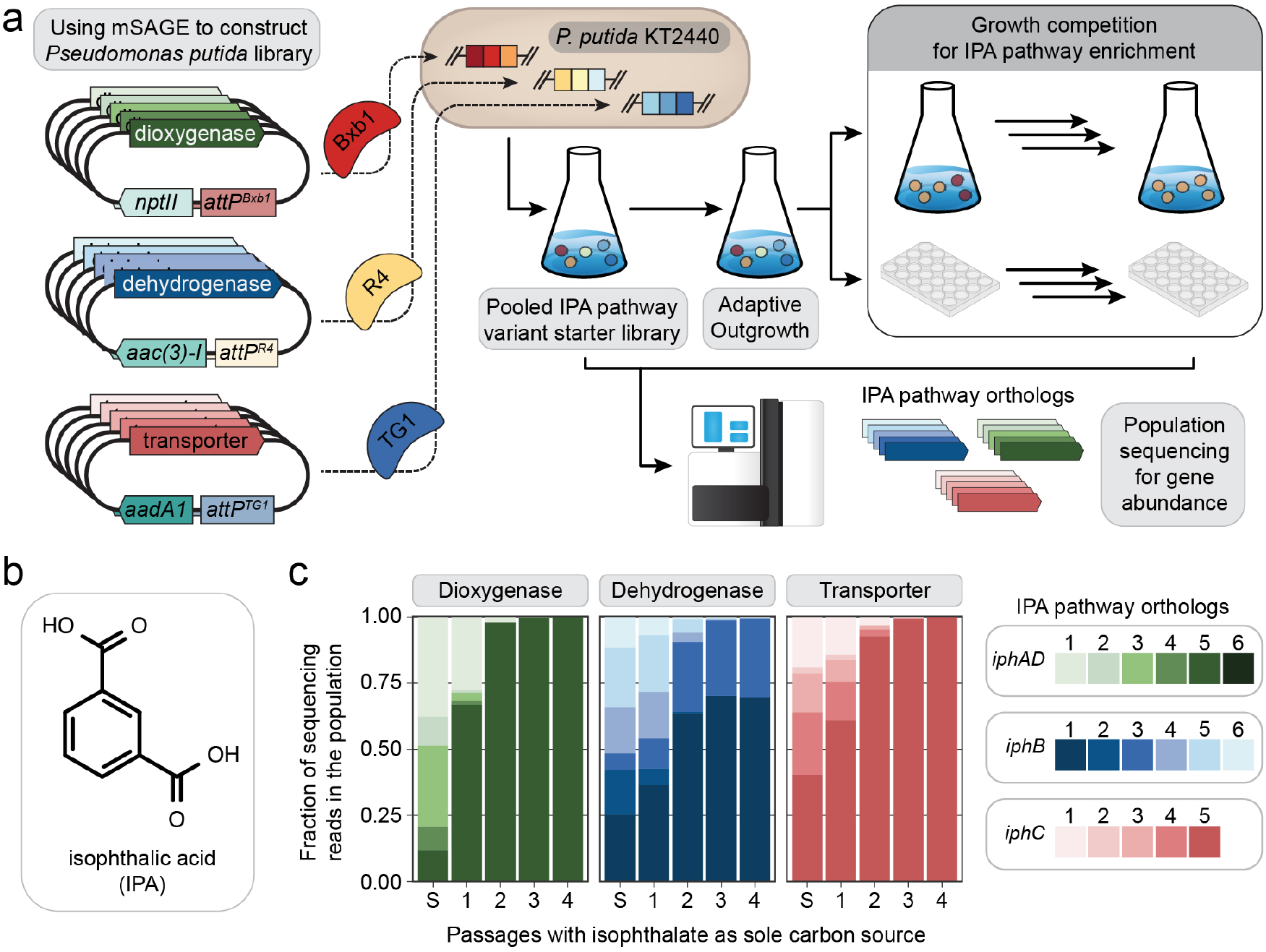
Application of mSAGE for non-native catabolic pathway optimization. **A.** Pooled mSAGE transformation of AG5577 with IPA pathway ortholog library followed by growth selection. **B.** Isophthalic acid chemical structure. **C.** Relative ortholog gene abundance during initial library construction (S) and subsequent outgrowth in minimal medium supplemented with IPA as a sole carbon source. Illumina PCR sequencing was used to quantify relative gene ortholog abundance at each stage of outgrowth. Ortholog protein sequences and sources are listed in Supplementary Table S3.

From the co-transformation, we created a pooled ∼32,000 member combinatorial library of IPA pathway components in a single transformation event, providing ∼175x coverage of the 180 possible combinations of dioxygenase, dehydrogenase, and transporter. Growth selection was utilized to identify ortholog combinations that enable the most rapid growth on IPA in *P. putida.* (Figure 2A). Direct transfer from rich medium into defined medium can cause stochastic lag phases with some aromatic carbon sources, so the pooled strain library was first pre-adapted to growth on aromatic compounds by cultivation in mineral medium with *p*-coumaric acid as the sole carbon source. Upon initial subculture into medium with IPA as the sole carbon and energy source, growth was observed after twenty-four hours and peaked within seventy-two hours, demonstrating the first transfer of IPA catabolism into a heterologous host. The pooled library was then sub-cultured in IPA for several additional passages to enrich for the fastest growing strains.

Samples were collected prior to and following each sub-culture to track enrichment of the best-performing pathway components (Figure 2A). Pooled PCR reactions and subsequent Illumina amplicon sequencing were performed to determine the relative ratio of each ortholog in the population. The original population was diverse, but a purifying selection for a subset of orthologs occurred rapidly (Figure 2B). Within two passages, *iphAD* and transporters were each dominated by a single variant, from *Acidovorax wautersii* and *Paraburkholderia tuberum*, respectively, and *iphB* resulted in two approximately equivalent variants from *Comamonas* E6 and *Maritimibacter alkaliphilus*. To validate that this combination of genes is best, we constructed ortholog combinations in which two of the best-performing components (e.g., the best dioxygenase and dehydrogenase) were combined with each of the individual third components (e.g., the transporters). Numerous ortholog combinations enabled growth on IPA. However, the most rapid growth on IPA occurred using the combination of orthologs that dominated the population in the competition assay (*iphAD* from *Acidovorax wautersii, iphB* from *Comamonas* E6, and MFS transporter from *Paraburkholderia tuberum*). This demonstrates that mSAGE and growth selection successfully enriched the population with the optimal ortholog combinations to achieve robust utilization of IPA as a growth substrate. Interestingly, the top combination featured components that each originated from different species.

Establishing an IPA catabolism pathway in *P. putida* expands the biological funnel required for efficient plastic waste bioconversion [16] and demonstrates the utility of mSAGE for generation of chromosomally integrated combinatorial libraries for metabolic engineering. This work emphasizes the importance of harnessing gene diversity in metabolic pathway development. In the future, the mSAGE toolkit can be applied to accelerate the solving of numerous scientific and engineering challenges that require integration of multiple heterologous DNA cargos. For example, co-optimization of dCas nuclease and guide RNA expression for CRISPR interference (CRISPRi), or for screening combinatorial libraries of guide RNAs for multiplexed CRISPRi. By allowing multiple rounds of chromosomal engineering, mSAGE enables the construction of deep combinatorial libraries while minimizing ancillary steps, such as marker removal or reintroduction of sites for recombination. The genetic parts that enable mSAGE, in particular the new recombinases, vector backbones, and multiplexed marker recycling with PhiC31, also benefit the base SAGE system, providing increased flexibility for single-cargo insertions. While the current mSAGE system allows for up to nine gene integrations, the development of additional recombinases [17] with high efficiency and minimal off-target integrations in *P. putida* would enable continued mSAGE cycles (Figure 1). Serine recombinases are functional in a wide array of bacteria. Thus, we expect mSAGE to allow for multiplexed chromosomal integration of multiple DNA fragments into many diverse bacteria, provided a limited number of gene expression parts are available in that host and transformation efficiency is sufficiently high. Thus, mSAGE expands upon the robust SAGE toolkit and will accelerate genetic engineering efforts to address societal challenges using *P. putida* and other organisms.

## Please see Supplemental Material below for materials and methods

## Acknowledgements

This work was authored by Oak Ridge National Laboratory, which is managed by UT-Battelle, LLC, for the U.S. Department of Energy (DOE) under contract DE-AC05-00OR22725. Funding was provided by the U.S. Department of Energy Office of Energy Efficiency and Renewable Energy Bioenergy Technologies Office (BETO) for the Agile BioFoundry. Funding was also provided in part by the US DOE, Office of Energy Efficiency and Renewable Energy, Advanced Materials and Manufacturing Technologies Office (AMMTO) and Bioenergy Technologies Office as part of the BOTTLE Consortium.

