## Supplemental Figures and Methods for "Combinatorial pathway engineering using multiplex Serine recombinase-Assisted Genome Engineering (mSAGE)"

**Table of Contents**

Supplemental Figure S1. Individual serine recombinase integration efficiency and accuracy.

Supplemental Figure S3. Collection of vectors used for mSAGE

Supplemental Figure S4. Multiplexed SAGE insertion of DNA into the *P. putida* chromosome

Supplemental Figure S5. Simultaneous removal of plasmid backbones using PhiC31 recombinase

Supplemental Figure S6 – Transport and initial catabolic pathway of isophthalic acid to the aromatic central metabolic intermediate protocatechuic acid

Supplemental methods

Supplemental Table S1. mSAGE plasmids and strain list

Supplemental Table S2. mSAGE oligonucleotide list

Supplemental Table S3 – IPA pathway ortholog sequences

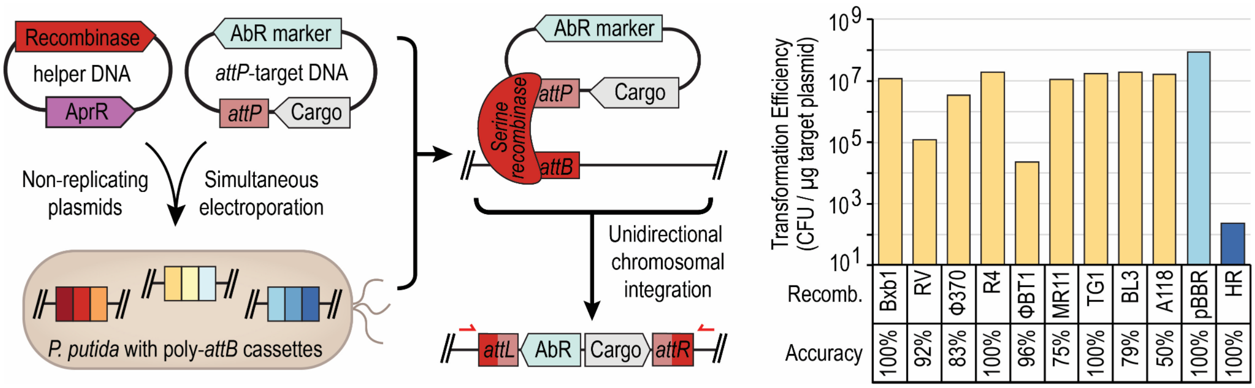


**Supplemental Figure S1.** Individual serine recombinase integration efficiency and accuracy. **A,** Individual SAGE *attP*-containing integrating *attP* plasmids were co-electroporated with the corresponding recombinase-expression plasmid to transform AG5577. This *P. putida* strain harbors *attB* sites at the following loci: *Bxb1, RV,* and *ΦC370* replace *hsdR (PP_4740)*; *R4, ΦBT1,* and *MR11* replace *ampC (PP_* 2876); and *TG1, BL3, and A118* are inserted into the intergenic region 3' of *fvpA* (*PP_4217).* **B.** Transformation efficiency (colony forming units (CFU) per microgram of cargo plasmid DNA) for each integrase, as well as the percent integration at the targeted *attB* site (accuracy). pBBR (light blue bar) is a replicating plasmid control and HR (dark blue bar) is a pK18mobsacB-based plasmid that inserts into the chromosome via homologous recombination. Data are the average and standard deviation of n ≥ 3 for transformation efficiency, and n ≥ 20 for accuracy.

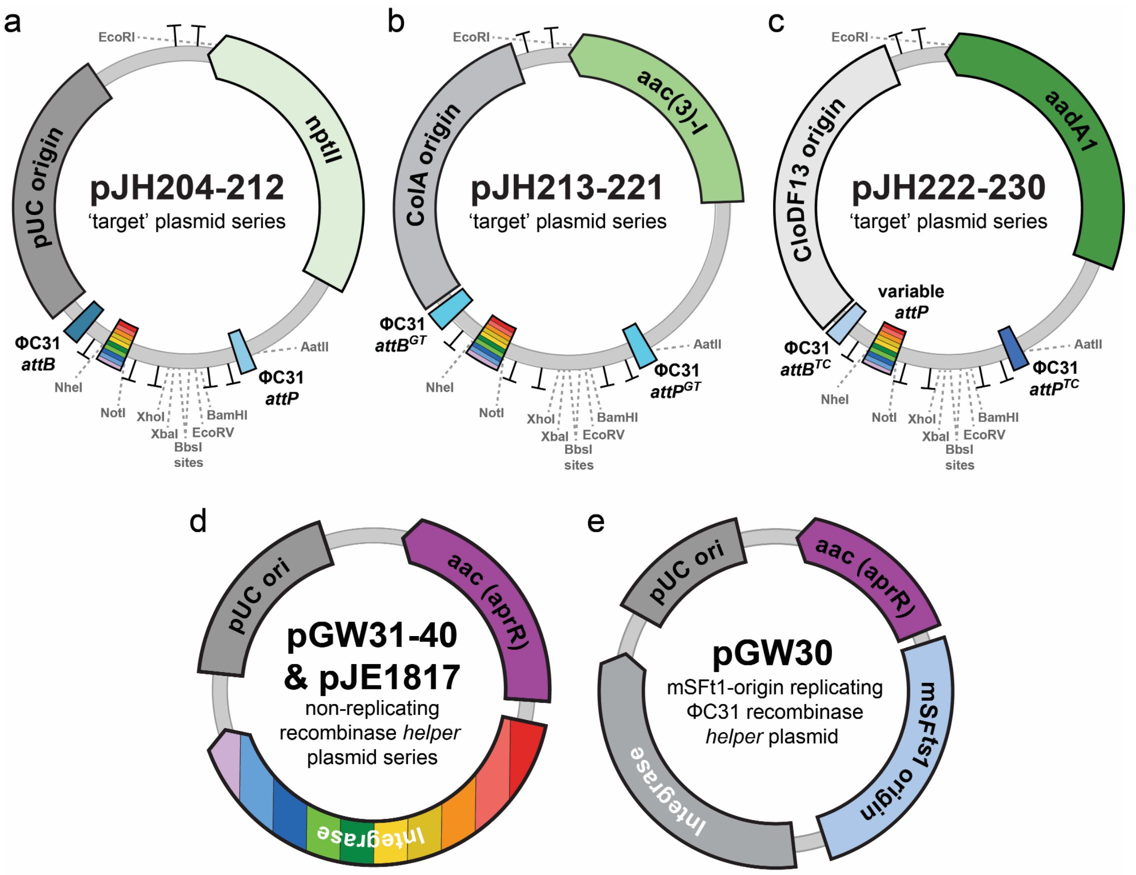


f

| Integrase | *nptII / pUC attP* target plasmid | *aac(3)-I / ColA attP* target plasmid | *aadA1 / CloDF13 attP* target plasmid | Non-replicating recombinase *helper* plasmid |
|---|---|---|---|---|
| Bxb1 | pJH204 | pJH213 | pJH222 | pGW31 |
| RV | pJH205 | pJH214 | pJH223 | pGW32 |
| Φ370 | pJH206 | pJH215 | pJH224 | pGW37 |
| R4 | pJH207 | pJH216 | pJH225 | pGW39 |
| ΦBT1 | pJH208 | pJH217 | pJH226 | pGW33 |
| MR11 | pJH209 | pJH218 | pJH227 | pGW36 |
| TG1 | pJH210 | pJH219 | pJH228 | pGW38 |
| BL3 | pJH211 | pJH220 | pJH229 | pGW40 |
| A118 | pJH212 | pJH221 | pJH230 | pGW35 |
| ΦC31 | --- | --- | pJE1817 | pGW34 |
| | | | | pJE1817 |

**Supplemental Figure S3.** Collection of vectors used for mSAGE. **A,** pJH204-212: cargo-free SAGE integrating vectors featuring ColE1 high-copy *E. coli* replicon, variable *attP* sites, and *nptII*. **B,** pJH213-221: cargo-free SAGE integrating vectors featuring ColA medium-copy *E. coli* replicon, variable *attP* sites, and *aac(3)-I*. **C,** pJH222-230: cargo-free SAGE integrating vectors featuring CloD13 low-copy *E. coli* replicon, variable *attP* sites, and *aadA1*. **D**, Non-replicating vectors pGW31-40 for transient recombinase expression during SAGE transformations. **E**, Temperature-sensitive vector pGW31 for SAGE cargo vector backbone removal (Supplemental Figure S2**)**. **F**, Table of mSAGE vectors and corresponding recombinase expressing vectors. AmpR, ampicillin resistance gene for selection in *E. coli*; *nptII*, kanamycin resistance gene for selection in *E. coli* and *P. putida*; *aac*, apramycin resistance gene for selection in *E. coli* and *P. putida*; *aadA1*, spectinomycin and streptomycin resistance gene for selection in *E. coli* and *P. putida*; *aac(3)-I* gentamycin resistance gene for selection in *E. coli* and *P. putida*; MCS, multiple cloning site. Additional mSAGE cargo vectors were constructed (pJH401-427, Supplementary Table S1) that are analogous to pJH204-230, but feature an *AmpR* marker for selection in *E. coli*, and ColA and CloDF13 replicons have been replaced by ColE1.

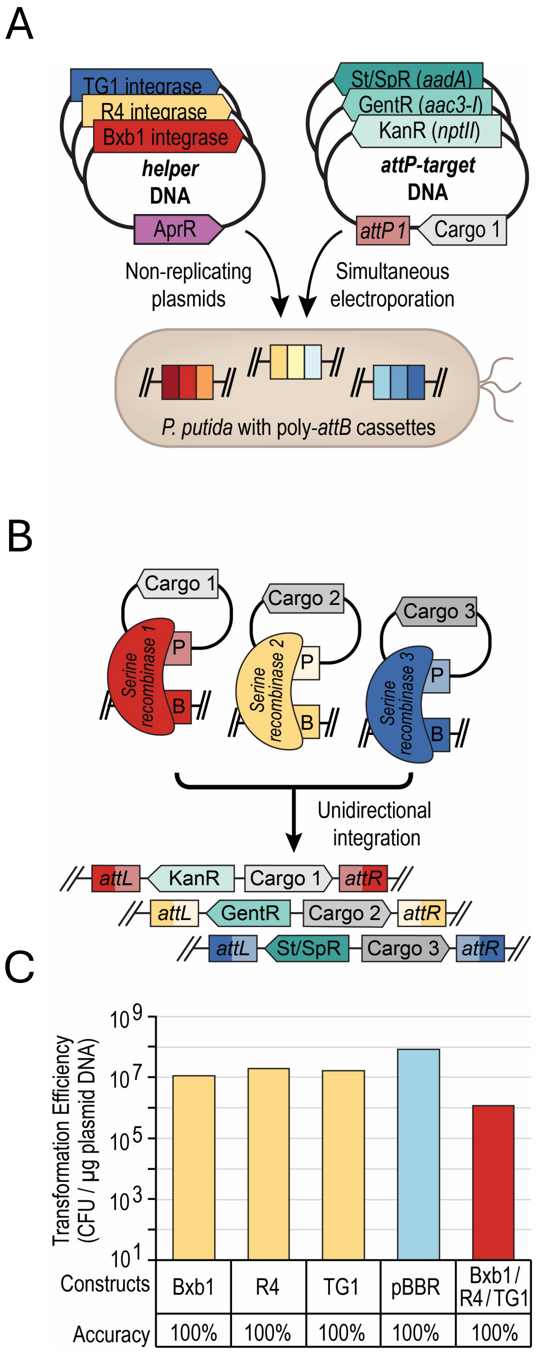


**Supplemental Figure S4.** Multiplexed SAGE insertion of DNA into the *P. putida* chromosome. **A,** mSAGE simultaneous integration of three SAGE cargo vectors in *P. putdia* KT2440. **B,** Transformation efficiency (CFU/µg cargo plasmid DNA) of single SAGE transformation, a replicating vector (pBBR), and mSAGE using *Bxb1*, *R4*, and *TG1* recombinase. Efficiency data are the average and standard deviation of n ≥ # for individual recombinases, and n ≥ # for triple insertion. Accuracy data are the average and standard deviation of n ≥ 20 for individual recombinases, and n ≥ 20 for triple insertion.

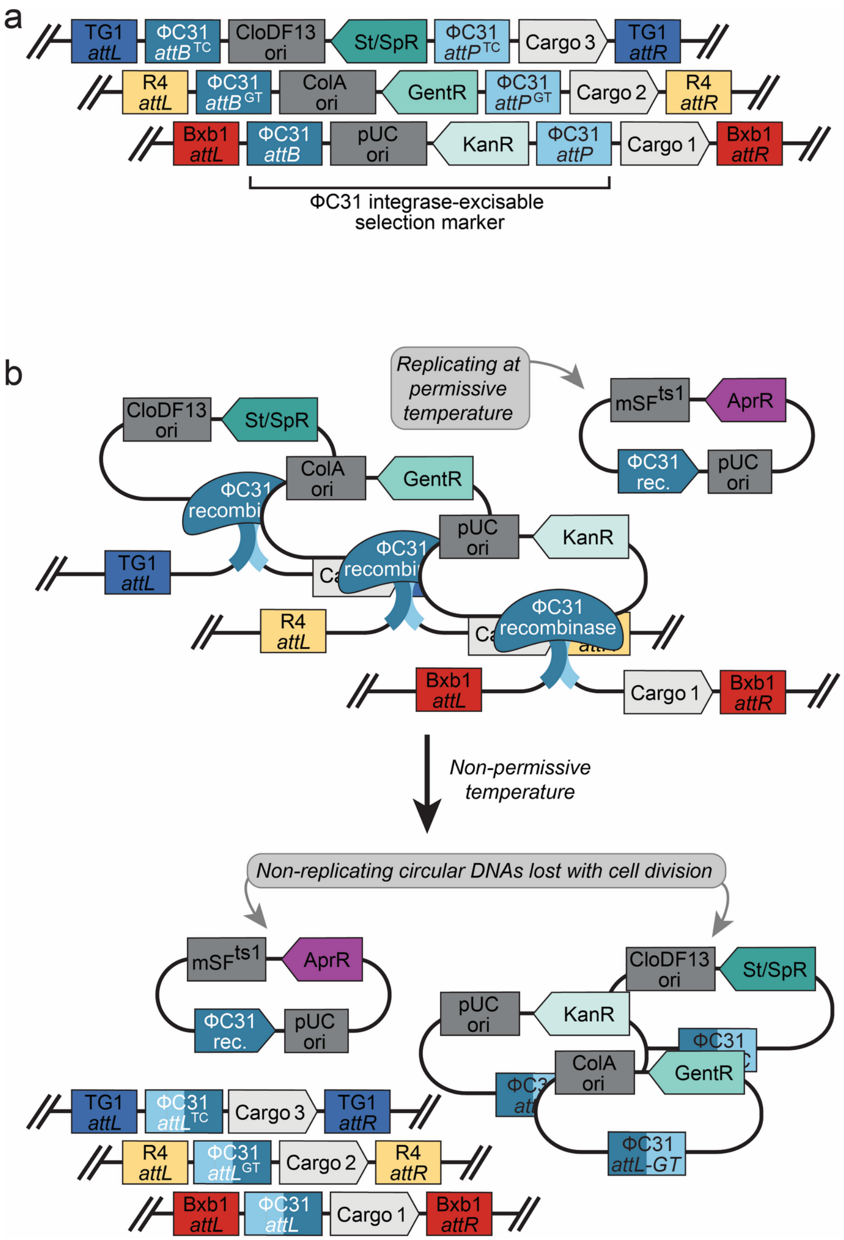


**Supplemental Figure S5.** Simultaneous removal of plasmid backbones using PhiC31 recombinase. **A,** Three SAGE cargo vectors that have been inserted into the chromosome of *P. putida*. **B,** Simultaneous removal of antibiotic resistance markers and *E. coli* replicons from *P. putida* KT2440 genome by *ΦC31* recombinase after mSAGE.

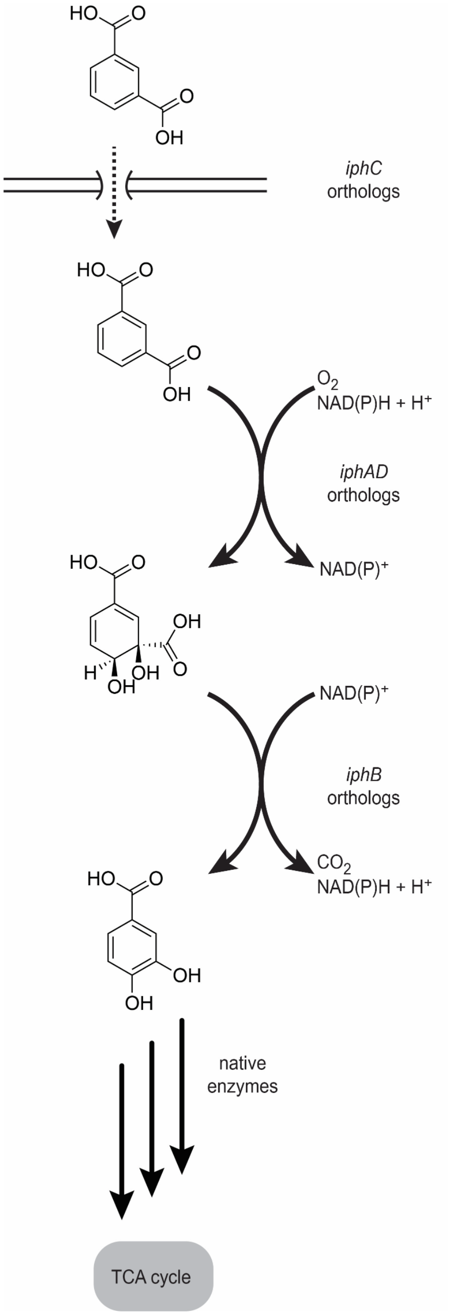


**Supplemental Figure S6.** Transport and initial catabolic pathway of isophthalic acid to the aromatic central metabolic intermediate protocatechuic acid. IphC; isophthalic acid transporter, IphAD; isophthalic acid dioxygenase and isophthalic acid dioxygenase reductase, IphB; 1,2-dihydro-1,2-dihydroxyisophthalic acid dehydrogenase.

**Supplementary Table S1: mSAGE Plasmids and Strain List**

**Note: All strains besides AG5577 are *E. coli* cloning strains harboring the listed plasmid.**

| Strain ID | Plasmid Name | Integrase *att*P | *E. coli* Origin | Resistance Marker | Plasmid Type | Source |
|---|---|---|---|---|---|---|
| **SAGE cargo vectors (empty)** | | | | | | |
| AG4513 | pJH0204 | Bxb1 | ColE1 | Kanamycin | Target/*att*P Plasmid | [7] |
| AG4514 | pJH0205 | RV | ColE1 | Kanamycin | Target/*att*P Plasmid | [7] |
| AG4515 | pJH0206 | phi370 | ColE1 | Kanamycin | Target/*att*P Plasmid | [7] |
| AG4516 | pJH0207 | R4 | ColE1 | Kanamycin | Target/*att*P Plasmid | [7] |
| AG4517 | pJH0208 | BT | ColE1 | Kanamycin | Target/*att*P Plasmid | [7] |
| AG4518 | pJH0209 | MR11 | ColE1 | Kanamycin | Target/*att*P Plasmid | [7] |
| AG4519 | pJH0210 | TG1 | ColE1 | Kanamycin | Target/*att*P Plasmid | [7] |
| AG4520 | pJH0211 | BL3 | ColE1 | Kanamycin | Target/*att*P Plasmid | [7] |
| AG4521 | pJH0212 | A118 | ColE1 | Kanamycin | Target/*att*P Plasmid | [7] |
| AG4522 | pJH0213 | Bxb1 | ColA | Gentamicin | Target/*att*P Plasmid | This work |
| AG4523 | pJH0214 | RV | ColA | Gentamicin | Target/*att*P Plasmid | This work |
| AG4524 | pJH0215 | phi370 | ColA | Gentamicin | Target/*att*P Plasmid | This work |
| AG4525 | pJH0216 | R4 | ColA | Gentamicin | Target/*att*P Plasmid | This work |
| AG4526 | pJH0217 | BT | ColA | Gentamicin | Target/*att*P Plasmid | This work |
| AG4527 | pJH0218 | MR11 | ColA | Gentamicin | Target/*att*P Plasmid | This work |
| AG4528 | pJH0219 | TG1 | ColA | Gentamicin | Target/*att*P Plasmid | This work |
| AG4529 | pJH0220 | BL3 | ColA | Gentamicin | Target/*att*P Plasmid | This work |
| AG4530 | pJH0221 | A118 | ColA | Gentamicin | Target/*att*P Plasmid | This work |
| AG4531 | pJH0222 | Bxb1 | CloDF13 | Spectinomycin/ Streptomycin | Target/*att*P Plasmid | This work |

| | | | | | | |
|---|---|---|---|---|---|---|
| AG4532 | pJH0223 | RV | CloDF13 | Spectinomycin/ Streptomycin | Target/*att*P Plasmid | This work |
| AG4533 | pJH0224 | phi370 | CloDF13 | Spectinomycin/ Streptomycin | Target/*att*P Plasmid | This work |
| AG4534 | pJH0225 | R4 | CloDF13 | Spectinomycin/ Streptomycin | Target/*att*P Plasmid | This work |
| AG4535 | pJH0226 | BT | CloDF13 | Spectinomycin/ Streptomycin | Target/*att*P Plasmid | This work |
| AG4536 | pJH0227 | MR11 | CloDF13 | Spectinomycin/ Streptomycin | Target/*att*P Plasmid | This work |
| AG4537 | pJH0228 | TG1 | CloDF13 | Spectinomycin/ Streptomycin | Target/*att*P Plasmid | This work |
| AG4538 | pJH0229 | BL3 | CloDF13 | Spectinomycin/Streptomycin | Target/*att*P Plasmid | This work |
| AG4539 | pJH0230 | A118 | CloDF13 | Spectinomycin/ Streptomycin | Target/*att*P Plasmid | This work |
| AG8142 | pJH401 | Bxb1 | ColE1 | Kanamycin/ Ampicillin | High Copy Target/*att*P Plasmid | This work |
| AG8143 | pJH402 | RV | ColE1 | Kanamycin/ Ampicillin | High Copy Target/*att*P Plasmid | This work |
| AG8144 | pJH403 | phi370 | ColE1 | Kanamycin/ Ampicillin | High Copy Target/*att*P Plasmid | This work |
| AG8145 | pJH404 | R4 | ColE1 | Kanamycin/Ampicillin | High Copy Target/*att*P Plasmid | This work |
| AG8146 | pJH405 | BT | ColE1 | Kanamycin/ Ampicillin | High Copy Target/*att*P Plasmid | This work |
| AG8147 | pJH406 | MR11 | ColE1 | Kanamycin/ Ampicillin | High Copy Target/*att*P Plasmid | This work |
| AG8148 | pJH407 | TG1 | ColE1 | Kanamycin/ Ampicillin | High Copy Target/*att*P Plasmid | This work |
| AG8149 | pJH408 | BL3 | ColE1 | Kanamycin/ Ampicillin | High Copy Target/*att*P Plasmid | This work |
| AG8150 | pJH409 | A118 | ColE1 | Kanamycin/Ampicillin | High Copy Target/*att*P Plasmid | This work |
| AG8151 | pJH410 | Bxb1 | ColE1 | Gentamicin/ Ampicillin | High Copy Target/*att*P Plasmid | This work |
| AG8152 | pJH411 | RV | ColE1 | Gentamicin/ Ampicillin | High Copy Target/*att*P Plasmid | This work |
| AG8153 | pJH412 | phi370 | ColE1 | Gentamicin/ Ampicillin | High Copy Target/*att*P Plasmid | This work |

| | | | | | | |
|---|---|---|---|---|---|---|
| AG8154 | pJH413 | R4 | ColE1 | Gentamicin/ Ampicillin | High Copy Target/*att*P Plasmid | This work |
| AG8155 | pJH414 | BT | ColE1 | Gentamicin/ Ampicillin | High Copy Target/*att*P Plasmid | This work |
| AG8156 | pJH415 | MR11 | ColE1 | Gentamicin/ Ampicillin | High Copy Target/*att*P Plasmid | This work |
| AG8157 | pJH416 | TG1 | ColE1 | Gentamicin/ Ampicillin | High Copy Target/*att*P Plasmid | This work |
| AG8158 | pJH417 | BL3 | ColE1 | Gentamicin/ Ampicillin | High Copy Target/*att*P Plasmid | This work |
| AG8159 | pJH418 | A118 | ColE1 | Gentamicin/ Ampicillin | High Copy Target/*att*P Plasmid | This work |
| AG8160 | pJH419 | Bxb1 | ColE1 | Spectinomycin/ Streptomycin/ Ampicillin | High Copy Target/*att*P Plasmid | This work |
| AG8161 | pJH420 | RV | ColE1 | Spectinomycin/ Streptomycin/ Ampicillin | High Copy Target/*att*P Plasmid | This work |
| AG8162 | pJH421 | phi370 | ColE1 | Spectinomycin/ Streptomycin/ Ampicillin | High Copy Target/*att*P Plasmid | This work |
| AG8163 | pJH422 | R4 | ColE1 | Spectinomycin/ Streptomycin/ Ampicillin | High Copy Target/*att*P Plasmid | This work |
| AG8164 | pJH423 | BT | ColE1 | Spectinomycin/ Streptomycin/ Ampicillin | High Copy Target/*att*P Plasmid | This work |
| AG8165 | pJH424 | MR11 | ColE1 | Spectinomycin/ Streptomycin/ Ampicillin | High Copy Target/*att*P Plasmid | This work |
| AG8166 | pJH425 | TG1 | ColE1 | Spectinomycin/ Streptomycin/ Ampicillin | High Copy Target/*att*P Plasmid | This work |
| AG8167 | pJH426 | BL3 | ColE1 | Spectinomycin/ Streptomycin/ Ampicillin | High Copy Target/*att*P Plasmid | This work |
| AG8168 | pJH427 | A118 | ColE1 | Spectinomycin/ Streptomycin/ Ampicillin | High Copy Target/*att*P Plasmid | This work |
| **Integrase expression plasmids** | | | | | | |
| AG3425 | pGW13 | Bxb1 | ColE1 | Apramycin | Integrase Expression TS Plasmid | [7] |
| AG3426 | pGW14 | BT | ColE1 | Apramycin | Integrase Expression TS Plasmid | [7] |

| | | | | | | |
|---|---|---|---|---|---|---|
| AG3429 | pGW17 | A118 | ColE1 | Apramycin | Integrase Expression TS Plasmid | [7] |
| AG3430 | pGW18 | MR11 | ColE1 | Apramycin | Integrase Expression TS Plasmid | [7] |
| AG3432 | pGW20 | phi370 | ColE1 | Apramycin | Integrase Expression TS Plasmid | [7] |
| AG3433 | pGW21 | RV | ColE1 | Apramycin | Integrase Expression TS Plasmid | [7] |
| AG3434 | pGW22 | TG1 | ColE1 | Apramycin | Integrase Expression TS Plasmid | [7] |
| AG3435 | pGW23 | R4 | ColE1 | Apramycin | Integrase Expression TS Plasmid | [7] |
| AG3436 | pGW24 | BL3 | ColE1 | Apramycin | Integrase Expression TS Plasmid | [7] |
| AG3438 | pGW30 | phiC31 | ColE1 | Apramycin | Backbone Excision Plasmid | [7] |
| AG3439 | pGW31 | Bxb1 | ColE1 | Apramycin | Integrase Expression Suicide Vector | [7] |
| AG3440 | pGW32 | RV | ColE1 | Apramycin | Integrase Expression Suicide Vector | [7] |
| AG3441 | pGW33 | BT | ColE1 | Apramycin | Integrase Expression Suicide Vector | [7] |
| AG3443 | pGW35 | A118 | ColE1 | Apramycin | Integrase Expression Suicide Vector | [7] |
| AG3444 | pGW36 | MR11 | ColE1 | Apramycin | Integrase Expression Suicide Vector | [7] |
| AG3445 | pGW37 | phi370 | ColE1 | Apramycin | Integrase Expression Suicide Vector | [7] |
| AG3446 | pGW38 | TG1 | ColE1 | Apramycin | Integrase Expression Suicide Vector | [7] |
| AG3447 | pGW39 | R4 | ColE1 | Apramycin | Integrase Expression Suicide Vector | [7] |
| AG3448 | pGW40 | BL3 | ColE1 | Apramycin | Integrase Expression Suicide Vector | [7] |

| | | | | | | |
|---|---|---|---|---|---|---|
| | **SAGE landing pad plasmids** | | | | | |
| AG4280 | pGW97 | ΔPP_4740::BxBI_RV_phi370_*att*B cassette | ColE1 | Kanamycin | Integrase Landing pad insertion plasmid | This work |
| AG4281 | pGW98 | ΔPP_2876::R4_phiBT1_MR11_*att*B cassette | ColE1 | Kanamycin | Integrase Landing pad insertion plasmid | This work |
| AG4282 | pGW99 | ΔPP_4217/4218 intergenic::TG1_BL3_A118_*att*B cassette | ColE1 | Kanamycin | Integrase Landing pad insertion plasmid | This work |
| | **IPA pathway ortholog plasmids** | | | | | |
| AG8046 | pJH465 | *Comamonas* sp. E6 *iph*AD | ColE1 | Kanamycin/ Ampicillin | Genetic cargo plasmid | This work |
| AG8047 | pJH466 | *Burkholderia* sp. CCGE10002 *iph*AD | ColE1 | Kanamycin/ Ampicillin | Genetic cargo plasmid | This work |
| AG8048 | pJH467 | *R. bacterium iph*AD | ColE1 | Kanamycin/ Ampicillin | Genetic cargo plasmid | This work |
| AG8049 | pJH468 | *P. tuberum iph*AD | ColE1 | Kanamycin/ Ampicillin | Genetic cargo plasmid | This work |
| AG8050 | pJH469 | *A. wautersii iph*AD | ColE1 | Kanamycin/ Ampicillin | Genetic cargo plasmid | This work |
| AG8051 | pJH470 | *C. testosteroni iph*AD | ColE1 | Kanamycin/ Ampicillin | Genetic cargo plasmid | This work |
| AG8052 | pJH471 | *Comamonas* sp. E6 *iph*B | ColE1 | Gentamicin/ Ampicillin | Genetic cargo plasmid | This work |
| AG8053 | pJH472 | *Burkholderia* sp. CCGE10002 *iphB* | ColE1 | Gentamicin/ Ampicillin | Genetic cargo plasmid | This work |
| AG8054 | pJH473 | *R. bacterium iphB* | ColE1 | Gentamicin/ Ampicillin | Genetic cargo plasmid | This work |
| AG8055 | pJH474 | *P. tuberum iphB* | ColE1 | Gentamicin/ Ampicillin | Genetic cargo plasmid | This work |
| AG8056 | pJH475 | *A. wautersii iphB* | ColE1 | Gentamicin/ Ampicillin | Genetic cargo plasmid | This work |

| | | | | | | |
|---|---|---|---|---|---|---|
| AG8057 | pJH476 | *C. testosteroni iphB* | ColE1 | Gentamicin/ Ampicillin | Genetic cargo plasmid | This work |
| AG8058 | pJH477 | *Comamonas* sp. E6 *iph*C | ColE1 | Spectinomycin/ Streptomycin/ Ampicillin | Genetic cargo plasmid | This work |
| AG8059 | pJH478 | *Burkholderia* sp. CCGE10002 *iphC* | ColE1 | Spectinomycin/ Streptomycin/ Ampicillin | Genetic cargo plasmid | This work |
| AG8060 | pJH479 | *A. wautersii iphC* | ColE1 | Spectinomycin/ Streptomycin/ Ampicillin | Genetic cargo plasmid | This work |
| AG8061 | pJH480 | *C. testosteroni iphC* | ColE1 | Spectinomycin/ Streptomycin/ Ampicillin | Genetic cargo plasmid | This work |
| AG8062 | pJH481 | *P. tuberum* MFS transporter | ColE1 | Spectinomycin/ Streptomycin/ Ampicillin | Genetic cargo plasmid | This work |

***Pseudomonas putida* KT2440 strains**

| **Strain ID** | **Plasmids used for strain construction** | **Genotype** | **Resistance Marker** | **Notes** | **Source** |
|---|---|---|---|---|---|

| | | | | | |
|---|---|---|---|---|---|
| AG5577 | pGW97-pGW99 | *Bxb1, RV, and ΦC370 attB sites replace hsdR (PP_4740); R4,ΦBT1, and MR11 attB sites replace ampC (PP_ 2876); and TG1, BL3, and A118 attB sites are inserted into the intergenic region 3' of fvpA (PP_4217). All attB landing pads are flanked by terminators.* | None | pGW97-pGW99 are homologous recombination vectors. Sucrose counterselection was used to ensure double-crossover and removal of *sacB*, *nptII*, and ColE1 origin. | This work |

**Supplementary Table S2: mSAGE Oligonucleotides**

| Primer Name | 5'-3' Sequence | Description | Purpose |
|---|---|---|---|
| oJH0716 | TCCCTTGTCCAGATAGCCCA | Forward primer targeting pJH204-212 backbones | Plasmid integration Screen PCR |
| oJH0717 | TGACGCTCAGTGGAACGAAA | Reverse primer targeting pJH204-212 backbones | Plasmid integration Screen PCR |
| oJH0718 | CTGTGTGAGCTCACAATTCC | Forward primer targeting pJH213-221 backbones | Plasmid integration Screen PCR |
| oJH0719 | TTCTAGGACGTTTCTGCGCA | Reverse primer targeting pJH213-221 backbones | Plasmid integration Screen PCR |
| oJH0720 | ACTGGGTTCGTGCCTTCATC | Forward primer targeting pJH222-230 backbones | Plasmid integration Screen PCR |

| | | | |
|---|---|---|---|
| oJH0721 | TTTCTACTGAACCGCGCATG | Reverse primer targeting pJH222-230 backbones | Plasmid integration Screen PCR |
| oJH0754 | CGACGATAGTGGCAGCATG | Upstream Flank targeting KT2440 ΔPP_4740 poly *att*B landing pad region | Plasmid integration Screen PCR |
| oJH0757 | AACGGCGTCAACCATTTTGT | Upstream Flank targeting ΔPP_2876 poly *att*B landing pad region | Plasmid integration Screen PCR |
| oJH0758 | GCAAGGCCAGGGTGAAATTG | Downstream Flank targeting ΔPP_2876 poly *att*B landing pad region | Plasmid integration Screen PCR |
| oJH0759 | CCGTACGACGTGTATGGTGA | Upstream Flank targeting ΔPP_4217/18 intergenic poly *att*B landing pad region | Plasmid integration Screen PCR |
| oJH0760 | CATGACCAGGGTGTCGCTTA | Downstream Flank targeting ΔPP_4217/18 intergenic poly *att*B landing pad region | Plasmid integration Screen PCR |
| oJH0761 | GCCACTCCCAAAATTCTGCC | Forward primer targeting BxbI and RV1 poly *att*B 1 region | Plasmid integration Screen PCR |
| oJH0762 | GGCAGAATTTTGGGAGTGGC | Reverse primer targeting BxbI and RV1 poly *att*B 1 region | Plasmid integration Screen PCR |
| oJH0763 | GTTCGCTCCACTAAAGTTGT | Forward primer targeting RV1 and phi370 poly *att*B 1 region | Plasmid integration Screen PCR |
| oJH0764 | ACAACTTTAGTGGAGCGAAC | Reverse primer targeting RV1 and phi370 poly *att*B 1 region | Plasmid integration Screen PCR |
| oJH0765 | ACTCCCCAATATGTGAGCA | Forward primer targeting R4 and phiBT1 poly *att*B 2 region | Plasmid integration Screen PCR |
| oJH0766 | CCCGTGCAACATCAGATGC | Reverse primer targeting R4 and phiBT1 poly *att*B 2 region | Plasmid integration Screen PCR |

| | | | |
|---|---|---|---|
| oJH0767 | AGAGTTGTCAGTTAGCTCGT | Forward primer targeting phiBT1 and MR11 poly *att*B 2 region | Plasmid integration Screen PCR |
| oJH0768 | ACCTGAACGAGCTAACTGAC | Reverse primer targeting phiBT1 and MR11 poly *att*B 2 region | Plasmid integration Screen PCR |
| oJH0769 | AACATGACTGCTGTCGGC | Forward primer targeting TG1 and BL3 poly *att*B 3 region | Plasmid integration Screen PCR |
| oJH0770 | GGCCCTTGTGCCGACAG | Reverse primer targeting TG1 and BL3 poly *att*B 3 region | Plasmid integration Screen PCR |
| oJH0771 | CGTCCAAGCGGATGCAATG | Forward primer targeting BL3 and A118 poly *att*B 3 region | Plasmid integration Screen PCR |
| oJH0772 | GGATCGCATTGCATCCGC | Reverse primer targeting BL3 and A118 poly *att*B 3 region | Plasmid integration Screen PCR |
| oJH0895 | CTGGAGTGGTGCACCCTG | Gene Target Screen forward primer-pJH465 | Plasmid integration Screen PCR |
| oJH0896 | GGCTGCCACTTGTACGGT | Gene Target Screen reverse primer-pJH465 | Plasmid integration Screen PCR |
| oJH0897 | ACCACGGCCTCAAGTTCG | Gene Target Screen forward primer-pJH466 | Plasmid integration Screen PCR |
| oJH0898 | CGGTATCGAGCACGGCAT | Gene Target Screen reverse primer-pJH466 | Plasmid integration Screen PCR |
| oJH0899 | GACCACATCTGCCCCCAC | Gene Target Screen forward primer-pJH467 | Plasmid integration Screen PCR |
| oJH0900 | GTAACGCCGATACCCCCG | Gene Target Screen reverse primer-pJH467 | Plasmid integration Screen PCR |
| oJH0901 | ACCACGGCCTGAAGTTCG | Gene Target Screen forward primer-pJH468 | Plasmid integration Screen PCR |
| oJH0904 | CCCAGCGCAGTACACCAA | Gene Target Screen reverse primer-pJH469 | Plasmid integration Screen PCR |

| | | | |
|---|---|---|---|
| oJH0905 | CGTCCTCGCCCCAAAGTT | Gene Target Screen forward primer-pJH470 | Plasmid integration Screen PCR |
| oJH0906 | ATGGCCCTGCAGCAAGTT | Gene Target Screen reverse primer-pJH470 | Plasmid integration Screen PCR |
| oJH0907 | GCAGGGATCGGGGCAG | Gene Target Screen forward primer-pJH471 | Plasmid integration Screen PCR |
| oJH0908 | TGAGGGTCGTACCGGTGA | Gene Target Screen reverse primer-pJH471 | Plasmid integration Screen PCR |
| oJH0909 | CAGGTGCTTCGAAGGGCA | Gene Target Screen forward primer-pJH472 | Plasmid integration Screen PCR |
| oJH0910 | AGAACACCTGCATGCGCT | Gene Target Screen reverse primer-pJH472 | Plasmid integration Screen PCR |
| oJH0911 | ACGAACTCAAGACCGCCG | Gene Target Screen forward primer-pJH473 | Plasmid integration Screen PCR |
| oJH0912 | CGTGTCTTGACGCTGGGT | Gene Target Screen reverse primer-pJH473 | Plasmid integration Screen PCR |
| oJH0913 | CCGTACTGCTCGTGGACG | Gene Target Screen forward primer-pJH474 | Plasmid integration Screen PCR |
| oJH0914 | CTCGAGGCCAGCCATACG | Gene Target Screen reverse primer-pJH474 | Plasmid integration Screen PCR |
| oJH0915 | GAGGCTATCCTGCAGGCG | Gene Target Screen forward primer-pJH475 | Plasmid integration Screen PCR |
| oJH0916 | ATTTCCTCCGGGCTTGCC | Gene Target Screen reverse primer-pJH475 | Plasmid integration Screen PCR |
| oJH0917 | TGTGTTCTGCGCTGAGGG | Gene Target Screen forward primer-pJH476 | Plasmid integration Screen PCR |
| oJH0918 | TCTGTCAGTGTGCTGCCG | Gene Target Screen reverse primer-pJH476 | Plasmid integration Screen PCR |

| | | | |
|---|---|---|---|
| oJH0919 | CCTCGGCCAACCATTCGT | Gene Target Screen forward primer-pJH477 | Plasmid integration Screen PCR |
| oJH0920 | GGTCGGGATGGTAGGGGT | Gene Target Screen reverse primer-pJH477 | Plasmid integration Screen PCR |
| oJH0921 | CGTAATCCAGGCCGCGAT | Gene Target Screen forward primer-pJH478 | Plasmid integration Screen PCR |
| oJH0922 | AGCGAACCCACCAGCATC | Gene Target Screen reverse primer-pJH478 | Plasmid integration Screen PCR |
| oJH0923 | TCGCACGTATGATCGCCC | Gene Target Screen forward primer-pJH479 | Plasmid integration Screen PCR |
| oJH0924 | AACTGCTTGGGGCTCGAC | Gene Target Screen reverse primer-pJH479 | Plasmid integration Screen PCR |
| oJH0925 | CGCATCGTGGTCCCCATT | Gene Target Screen forward primer-pJH480 | Plasmid integration Screen PCR |
| oJH0926 | GGGTGCTGGCAATCGTCT | Gene Target Screen reverse primer-pJH480 | Plasmid integration Screen PCR |
| oJH0927 | TGGGCATCTCCGTCTCGA | Gene Target Screen forward primer-pJH481 | Plasmid integration Screen PCR |
| oJH0928 | GATCGGCGCGACCAAGTA | Gene Target Screen reverse primer-pJH481 | Plasmid integration Screen PCR |
| oJH0966 | GTCCGGATCCCTATGGAGGTCAGG | Froward primer for IPA strain sequence screen | Pooled PCR for Population Tracking |
| oJH0967 | GTCGGCCTCAGCGATCTCGC | Reverse primer for IPA strain seq screen-pJH465 | Pooled PCR for Population Tracking |
| oJH0968 | GGCTCGGCGATTTCGCTCGC | Reverse primer for IPA strain seq screen-pJH466 | Pooled PCR for Population Tracking |
| oJH0969 | CCTTCTCCAGGTCGCTCGACG | Reverse primer for IPA strain seq screen-pJH467 | Pooled PCR for Population Tracking |

| | | | |
|---|---|---|---|
| oJH0970 | CTGGCTCCTCAATTTCGGACG | Reverse primer for IPA strain seq screen-pJH468 | Pooled PCR for Population Tracking |
| oJH0971 | CGCATCCGGGATTTCGCTGC | Reverse primer for IPA strain seq screen-pJH469 | Pooled PCR for Population Tracking |
| oJH0972 | GGCCTCCGCGATTTCGGACG | Reverse primer for IPA strain seq screen-pJH470 | Pooled PCR for Population Tracking |
| oJH0973 | GTCGACCATCAGCACGGCCG | Reverse primer for IPA strain seq screen-pJH471 | Pooled PCR for Population Tracking |
| oJH0974 | GCGCTACCAGGGTGACCTGC | Reverse primer for IPA strain seq screen-pJH472 | Pooled PCR for Population Tracking |
| oJH0975 | ACCAACACGACAGCGGCCCC | Reverse primer for IPA strain seq screen-pJH473 | Pooled PCR for Population Tracking |
| oJH0976 | ACGAGCAGTACGGCAGCGCC | Reverse primer for IPA strain seq screen-pJH474 | Pooled PCR for Population Tracking |
| oJH0977 | GACCATCAGCACTGCCGCGC | Reverse primer for IPA strain seq screen-pJH475 | Pooled PCR for Population Tracking |
| oJH0978 | CCAGCACGACAGCCGCGCC | Reverse primer for IPA strain seq screen-pJH476 | Pooled PCR for Population Tracking |
| oJH0979 | CGGACCGGTTTGGTCGGGCG | Reverse primer for IPA strain seq screen-pJH477 | Pooled PCR for Population Tracking |
| oJH0980 | GAAGAGGGCAACCCACCGG | Reverse primer for IPA strain seq screen-pJH478 | Pooled PCR for Population Tracking |
| oJH0981 | TACGGACCGGCTTCGTCGGC | Reverse primer for IPA strain seq screen-pJH479 | Pooled PCR for Population Tracking |
| oJH0982 | CGTACCGGTTTTGTTGGGCGCC | Reverse primer for IPA strain seq screen-pJH480 | Pooled PCR for Population Tracking |
| oJH0983 | CAGGGCAACCCAGCGGTACG | Reverse primer for IPA strain seq screen-pJH481 | Pooled PCR for Population Tracking |

**Supplementary Table 3: IPA pathway ortholog sequences**

**Note: (#) corresponds to ortholog # in Figure 2C.**

| **Name** | **Source Organism** | **Sequence** |
|---|---|---|
| iphA (1) | *Comamonas* sp. E6 | MNKEMSETLTRVGPNTRMGNLLRRYWVPALMSSEIAEADGPQVRVQLLGEKLLAFRNTDGKACL ISEFCSHRGVSLYFGRNEENGIRCAYHGVKFDGDGQCVDVPSSPQSCARMHIKGYPCVERGGIVW TYMGPEEHKPSPPELEWCTLPPEHVFVSKRLQYSNWLQAMEGGIDTAHVSYVHRFEVDTDPMHQ GVKALDYIKADGNVKFEIEQTPFGLSLFGRRNGEPDSYYWRITQWLFPWFTLIAPFGQHALGGHV WVPIDDHNCWAWSINWQPDQPLTEEERTSMEEGKGIHVEYEAPGSFIPKANRNNDYGMDRVAQR EERSYSGIFGFSAQDYSLQESMGPIQDHAAERLLPTDKAIVMARRMLNEAALGLEQGETPPALDAS EQHVRPAGVLLPRDQDPVAWAREELADATKKPVFSL* |
| iphA (2) | *Burkholderia* sp. CCGE10002 | MDKNMSETLVRTGPGTAMGNLMRRYWVPVLLASEIAEPDCPPVRVQILGEKLLAFRDSEGQPALI DEFCSHRGVSLYFGRNEENGIRCSYHGLKFDRNGNCVEVPSAPQACKHMGITAYPCIERAGIVWA YMGPKDRQPAPPDLEWCNLPDSHVFVSKRLQESNYLQAMEGGIDTSHVSYVHRYEVDDDPMHQ GTKALDYIKADGNVIFEIEKQPFGLTLFGRRNGEPDSYYWRVTQWLFPWYTLIPPFGDHSLAGHV WVPIDDHSCWAWSINFRPDRPLDEQELADLNAGKGIHCEYEEGGSFRPKANKDNDYLIDRKAQKE KRAYSGVFGFAMQDASLQESMGPIQDHSKEKLLPTDRAIVMARRMMYEAATALVPDTDPPAIDA DQQRVRAAGVLLPRDQKPQEWAVIHLHDGKDQPIYTI* |
| iphA (3) | *Maritimibacter alkaliphilus* HTCC2654 | MLSHEDNETLVRVGPGTTMGDMMRLYWLPFMASSDLEKDGQPQTVKLLDETLIVFRDSE GRVGLVDHICPHRGAPLVFGRNEDCGLRCVYHGWKFDVDGNVADMPAEPPRSRLKDRV KIKSYPCVERGGVVWTYMGDEPDESRPALPSFEWNMVPEENVVVTFRVQECNWLQALE GEIDSAHAPILHGRIDDGGSINQWVAKRDLRPTFECMRQDFGMSIASRRVLDDDTLYWRV |

| | | |
|---|---|---|
| | | NQFVMPFFSLVPPQSNEFYELSGHAWVPIDDENTLCIMFSYRPDEPLHPKSRKVFLDGHNG<br>RETGHPSREGFDDQGAQVPFGRYVSKYRRETGWLFDNEAQKTTWFSGLPGLWVQDAAC<br>QSGVLRVYDRTREHLCTSDTGIAMTRRMLLETAHAFRDSGKKPDRFDDPDLYLVRAVSL<br>KLPKDLPWAEAGKAPMTAKVGEGLGYEL* |
| iphA (4) | *Paraburkholder ia tuberum* environmental isolate | MDKNMSETLVRTGPGTAMGNLMRRYWVPVLLSSEIEEPDCPPVRVQILGEKLLAFRDSEGQPALI<br>DEFCSHRGVSLFFGRNEENGIRCSYHGLKFDRNGNCVEVPSAPQACKHMGITAYPCIERAGIVWTY<br>MGPKDRQPAPPDLEWVSLPASHVFVSKRLQESNYLQAMEGGIDTSHVSYVHRYEVDDDPMHQGT<br>KALDYIKADGNVIFEIEKQDFGLTLFGRRNGEPDSYYWRVTQWLFPWYTLIPPFGDHSLAGHVWV<br>PIDDHSCWAWSVNFRPDRPLDEQELADLNAGKGIHCEYEEGGKFRPKANKDNDYLIDRQAQKDK<br>RAYSGVFGFAMQDASLQESMGPIQDHSKEKLLPTDRAIVMARRMLYEAATALVPDTDPPAIDANQ<br>QRVRAAGVLLPRDQKPQEWAVIHLHDGKDQPIYSI* |
| iphA (5) | *Acidovorax wautersii* environmental isolate | MNKEMSETLTRVGPGTRMGNLMRRYWVPALACSEIPDADGPQVRVQLLGEKLLAFRNSDGQACL<br>IGEFCSHRGVSLYFGRNEQNGIRCAYHGVKFDGMGQCVDVPSSPQACSRMHIKGYPCVERGGIVW<br>AYMGPADQQPAPPELEWCTLPPEHVFVSKRLQYSNWLQAMEGGIDTAHVSYVHRFEVDTDPMH<br>QGVKALDYIKADGNVKFEIEQTPFGLSLFGRRNGEEDSYYWRITQYLFPWFTLIAPFGEHALGGHV<br>WVPIDDHHCWAWSINWQPGQPLTTEERTAMEEGKGIHVEYEAPGSFIPKANRDNDYGMDRVAQ<br>KEERSYSGIFGFSAQDYSLQESMGSIQDHEAEKLLPTDKAIVMARRMLHEAALGLEQGQTPPALD<br>AREQHVRPAGVLLPRDQDPVAWAREELADATKKPVFSL* |
| iphA (6) | *Comamonas testosteroni* environmental isolate | MNKEMSETLTRVGPNTRMGNLLRRYWVPALMSSEIAEADGPQVRVQLLGEKLLAFRNTDGKACL<br>ISEFCSHRGVSLYFGRNEENGIRCAYHGVKFDGDGQCVDVPSSPQSCARMHIKGYPCVERGGIVW<br>TYMGPEEHKPSPPELEWCTLPPEHVFVSKRLQYSNWLQAMEGGIDTAHVSYVHRFEVDTDPMHQ<br>GVKALDYIKADGNVKFEIEQTPFGLSLFGRRNGEPDSYYWRITQWLFPWFTLIAPFGNHALGGHV |

| | | |
|---|---|---|
| | | WVPIDDHNCWAWSINWQPDQPLTKEERQSMEEGKGIHVEYEEPGSFIPKANRNNDYGMDRVAQR<br>EERSYSGIFGFSAQDYSLQESMGPIQDHAAERLLPTDKAIVMARRMLNEAALGLEQGETPPALDAS<br>EQHVRPAGVLLPRDQDPVAWAREELADATKKPVFSL* |
| iphB (1) | *Comamonas* sp. E6 | MSESRMAGRTALITGAGAGIGAAASHLFCQEGAAVLMVDANAEALERTREAILQAVPGARLACA<br>TADVSDEAAAAAAVGQCVQQWGGLDTLVNNAAMRNYSAAADATAAEWQAMVGVNLVGMSN<br>YCRAALPALRQSGTGSIVNVSSCYAVTGRKGMALYDATKAAQLAYTRSLAFEEAAHGVRANAVC<br>PGSTLTDFHVGRARNAGKSVEQLRTERKDTSLIGRWASPEEIAWPILWLASSEASFITGTTLMVDG<br>GLHIM* |
| iphB (2) | *Burkholderia* sp. CCGE10002 | MKQRKRLEDKVALITGGGGGIGAATARVFCAEGAAVVLVDANQEALARVADELKTADPSARVET<br>FAADVSNDADAMRAVQLAADSFGRLDVLVNNAAMRNYSALADATPAEWQAMVSVNLVGTSNY<br>CSAALPFLRRAGRASIVNVSSCYAVTGRKGMGLYDATKAGMLAMTRTLAFEETANGVRVNAVCP<br>GSTLTEFHVNRAIASGKSVEVLKTQRQDTSLIGRWASPEEIAWPILWFASDEASYITGTTLMVDGG<br>LSAM* |
| iphB (3) | *Maritimibacter alkaliphilus* HTCC2654 | MDLNLSGLNALVTGASKGIGLATARTLAQEGVQVTLVARDCDRLADAAAQIEGDTGLRP<br>DVVAADLATREGVERVAARETPVDILVNNAGAIPPGDLASIDEDRWRAAWDLKVFGYIN<br>MTRALAPRIAERGGVIVNVIGAGGEALSADYICGAVGNAALMAFTRAYAKQFNSMGGRI<br>VGLNPGLVATERMQVFLKSRAAAELGDEGRADELTRGLPYGRAADPQEVADAVAFLAS<br>PRSGYTNGTILSLHGGG* |
| iphB (4) | *Paraburkholderia tuberum* | MQQRKRLEDKVALITGGGGGIGAATARVFCAQGAAVVLVDANPEALARVADELKTADPSARVET<br>FAADVSHEGDAARAIQLAADSFGKLDVLVNNAAMRNYSALADATPAEWQAMVSVNLIGTSNYC<br>RAALPFLRRAGRASIVNVSSCYAVTGRKGMGLYDTTKAGMLAMTRTLAFEEAAHGVRANAVCP |

| | | |
|---|---|---|
| | environmental isolate | GSTLTDFHVNRAVAAGRSVEVLKTQRQDTSLIGRWASPEEIAWPILWFASDEASYITGTTLMVDG<br>GLSAM* |
| iphB (5) | *Acidovorax wautersii* environmental isolate | MGEQRLAGRTALITGAGAGIGAAAAHLFCREGAAVLLVDANGDALERTRQAIADAVPGARLACA<br>TADVSDEAAALAAVQQCTGAWGGLDILVNNAAMRNYSAAADATAAEWQAMVGVNLVGMSHY<br>CRAALPALRQSGAGSIVNVSSCYAVTGRKGMALYDATKAAQLAYTRTLAFEEAPHGVRANAVCP<br>GSTLTDFHVGRAQTAGKSVDQLRTERKDSSLIGRWASPEEIAWPIVWLASSEASFITGTTLMVDGG<br>LHIM* |
| iphB (6) | *Comamonas testosteroni* environmental isolate | MSESRMAGRTALITGAGAGIGAAASHLFCKEGAAVLMVDANAEALERTREAILQAVPGARLACV<br>MADVSDESAAAAAVGQCVQQWGGLDTLVNNAAMRNYSAAADATAAEWQAMVGVNLVGMSN<br>YCRAALPALRQSGTGSIVNVSSCYAVTGRKGMALYDATKAAQLAYTRSLAFEEAAHGVRANAVC<br>PGSTLTDFHVGRARNAGKSVEQLRTERKDTSLIGRWASPEEIAWPIFWLASREASFITGTTLMVDG<br>GLHIM* |
| iphC (1) | *Comamonas* sp. E6 | MKFPMTLKAALVAGACAATAAWGQAAAAPEWRPTKPVRIVVPITGSTNDVLARLIAPKLQEALG<br>QPFVVENKPGAGGNIGAYEVSRSVPDGHTLLIGYNGPLAINVTLFDKMPYDPLKDLAPITLAVKSP<br>QYLVVNPKAEFKDVKDFIAKAKANPSKYSYGSVAMGSASHLTMEMMKSAAGFQMTHVPYKGA<br>GPAVTDLIAGNVQSGFFVPGNVQGFVKEGRLKLLASTGAKRFPSTPTIPTLAESGLKDFEATSWIGL<br>LAPAGTPPAVINTYHQAMVRILNSPDVRKHLDEMEFETIASTPKQFSDWIATEIGRWGKVIKATNA<br>KAD* |
| iphC (2) | *Burkholderia* sp. CCGE10002 | MEKNLASSQGMSASVWSSNGAREASSADTNPYRWVALFIVWAAFLLSYVVRVAWSTVAAPVGA<br>SLGISVSMLGAFVTAFYAGYVLANVVGGMLTDLLGGRAMLTLALLPLGVLTFTFGYTHSLAFGIVI<br>QAAMGFAAGADYSAGMKIIPAWFTRDRGRAMGLYTTATSLAVVIANAVVPSFSARHGWSNAFH<br>MLGIVTFAWGIVTLLLLKNRPSNEAKPARNSLQDMLGLLRNRNLIALSIAGCGGLWATVGFAAWG |

| | | |
|---|---|---|
| | | NALMTRQYGIAPIVAGSIVASFGIGAVIAKPTLGWISDLPGVSRRMMSIGCLIAFTIALLVFGYCSTA HQFYLVAPILGAFSFGYLPVLMAQVSDASGKRLAGASAGWTNAIWQSGSAVSPMLVGSLYGASH SFLLALITLAIGPAIAVVAMFFVNPQIARE* |
| iphC (3) | *Acidovorax wautersii* environmental isolate | MKFPMKLAAGLALLACASSTAWAQAAASDWRPTKPVRIVVPITGSTNDVLARMIAPKLQEALGQ PFFVDNKPGAGGNIGAAEVARSAPDGHTLLIGYNGPLAINVTLFDKMPYDPQKDLTPITLAVKSPQ YLVVNPDTGVTDVKDFIAKAKADPKKFAYGSVAMGSASHLTMEMMKSAAGFQATHVPYRGAGP AVTDLIAGNIQAGFFVPGNVQGFVKEGRLKLLASTGPKRFPSTPDIPTLAESGLKDFEATSWIGLLA PANTPPAIVSSYHQAMVRILNSPDITKRLQEMEFEVVASSPKQFSDWIGTEITRWGKVIKATGAKAE * |

**Table S3 Continued**

| Name | Source Organism | Sequence |
|---|---|---|
| iphC (4) | *Comamonas testosteroni* environmental isolate | MKFPMTLKAALVAGACAATAAWGQAAAAPEWRPTKPVRIVVPITGSTNDVLARLIAPKLQEALGQPFVVE NKPGAGGNIGAYEVSRSVPDGHTLLIGYNGPLAINVTLFDKMPYDPLKDLAPITLAVKSPQYLVVNPKAGF KDVKDFIAKAKANPSKYSYGSVAMGSASHLTMEMMKSAAGFQMTHVPYKGAGPAVTDLIAGNVQSGFFV PGNVQGFVKEGRLKLLASTGAKRFPSTPTIPTLAESGLKDFEATSWIGLLAPAGTPPAVINTYHQAMVRILNS PDVRKHLDEMEFETIASTPKQFNDWIATEIGRWGKVIKATNAKAD* |
| MFS Transporter (5) | *Paraburkholderia tuberum* environmental isolate | MEKNLASSEGMAAPVWSANDAPAASSADTSPYRWVALFIVWAAFLLSYVVRVAWSTVAAPVGASLGISV SMLGAFVTAFYAGYVLANVVGGMLTDLLGGRAMLTLALLPLGVLTFTFGYTHSLAFGIVIQAAMGFAAGA DYSAGMKIIPAWFTRDRGRAMGLYTTATSLAVVIANALVPSFSARHGWSNAFHMLGIVTFGWGIVTLLLLR NRPSNEAKPARNSVQEMLGLLRNRNLIALSIAGCGGLWATVGFGAWGNALMTRQYGIAPVVAGSIVASFGI |

| | | |
|---|---|---|
| | | GAVIAKPTLGWISDLPGVSRRMMSIGCLTAFAIALLVFGYCSTVREFYLVAPILGAFSFGYLPVLMAQVSDA<br>SGKRLAGASAGWTNAIWQSGSAVSPMVVGSLYGASHSFMLALITLAIGPAVAVVAMFFVNPHIARE* |
| iphD (1) | *Comamonas* sp. E6 | MASTYLQARVHQMRYEAAGTLSVELRPLVVAEEFAQPVQAGAHIDLHLADGLIRSYSLINPGERHRYVVA<br>VSLDPASRGGSRFVHEKLRVGQAIQIGGPRNHFPLDETASHSVLVAGGIGITPVLSMLRKLHALGRTAHLIY<br>CASSRENAAFVPEIEAIAAQAGGRVTVDWHFKDEKGVRADLYSLLQGHAEGAHFYACGPLVFLASYEDSC<br>QKLGLAHVHLERFAAAPLAAPQTPEVGYAVELRRTGKTVQVAAGTSLLDTLINAGMNPEYSCREGVCGAC<br>EVRVISGDVDHRDQILSEQERAANKSMMICVSGCRSGNLVLDC* |
| iphD (2) | *Burkholderia* sp. CCGE10002 | MNMTDSTLMMRVAARRDEADGIAGFEFVDADGRELPPFEAGAHIDVYVPGGPVRQYSLCNAPHERHRYQI<br>AVLRDANSRGGSQRMHDAVNEGDAIHIGVPRNHFPLARHDAKPLLLAGGIGVTPILCMAEQLAAKGAAFD<br>MHYCARSKSRAAFVERIAASSWADNVQYHFDDEHGMLDLNALLTGGADRHLYVCGPQGFMNAVLDTAR<br>SLGWSDDRLHYEYFAAAQPSGDGESFDVRLARSGRVVSIAADCTVTQALAAAGVDVPVSCEQGICGTCITR<br>VLDGEPDHRDLYLSPEEQARNDQFLPCCSRAKSRVLVLDL* |
| iphD (3) | *Maritimibacter alkaliphilus* HTCC2654 | MSKGADIPVRVRQMRYEADTILSVEFETLDGRDLPVAAPGSHIDVALGKDFRRSYSLTREITGGPSCTVAIH<br>RDPKSKGGSAYVHETLRVGDKTVISRPKNNFPLDENADLSVLIAGGIGVTPVLCMIRRLVATGKAWKLHYA<br>ARSRSAAAFRDELDALEQASDGKGKLHYHLDDEQNGALVDIGGILTANPTAHFYCCGPEAMLAAYERAAR<br>GVPRDQVHVEYFSSSEEVARDGGFEVVLDRSGKTIVVEPGQTILDALIANGVHVPFSCAEGTCGTCETDVIE<br>GRPDHRDIILTDEERAESKTMMICCSGSKSARLVLDI* |
| iphD (4) | *Paraburkholderia tuberum* environmental isolate | MNMTDSTLMVRVAARRDEADDIAGFEFVDVDGRELPPFEAGAHIDVYVPGGPVRQYSLCNAPGERHRYQI<br>AVLRDAGSRGGSQRMHDAVNPGDAIRIGVPRNHFPLAPHDAQPLLLAGGIGVTPILCMAEQLAATGAAFE<br>MHYCARSKSRAAFVERIAASPWAARVQYHFDDEHGVLDLRALLAGVSAGRHLYVCGPQGFMNAVLDTA<br>RSLGWSDDRLHYEYFAAAQPAGDGGSFDVRLARSGRVVPVAADCTVTQALAAAGVDVPVSCEQGICGTCI<br>TRVLEGEPDHRDLFLSPEEQARNDQFLPCCSRAKSRVLVLDL* |

| | | |
|---|---|---|
| iphD (5) | *Acidovorax wautersii* environmental isolate | MAVMTTLQARIHQLRYEAAGTTSVELRPLPPATQFAQPVEAGAHIDLHLADDLTRSYSLTNPGEAHRYVVA<br>VARDPASRGGSRFVHESLRVGQVITIGGPRNHFALDESAPHSVLVAGGIGITPVYAMLQRLAVLGRTAHLV<br>YCAGSRAGAAFVADIEALAAASAGSITVDWYFKDERGTRADLPRLLAGHPEGTHFYACGPLPLLDGYEQA<br>CESLGLAHVHLERFAAAPLAPSQTPSQGYDVELRKSGKTVHVAPGIALLDALLDAGMNPDYSCREGVCGA<br>CEMKVLCGEVDHRDLILSKQDQAANRSMMVCVSGCKSGSLVLDF* |
| iphD (6) | *Comamonas testosteroni* environmental isolate | MASTYLQARVHQMRYEAAGTLSVELRPLVVAEEFAQPVQAGAHIDLHLADGLIRSYSLINPGESHRYVVA<br>VSLDPASRGGSRFVHQRLRVGDVIQIGGPRNHFPLVETAPHSVLVAGGIGITPVLAMLRRLDALGRTAHLIY<br>CASSRASAAFVPEIEAIAAQVGGRVTVDWHFKEEKGVRADLHNLLQGHAEGAHFYACGPLAFLDSYEDSC<br>GKLGLAHVHLERFAAAPLAAPRTPEVGYAVELRRTGRTVQVAAGTSLLDTLINAGMNPEYSCREGVCGA<br>CEVRVISGDVDHRDQILSEQERAANKSMMICVSGCRSGNLVLDC* |

**Supplemental materials and methods:**

### Culture conditions

Generally, strain propagation of *P. putida* or *E. coli* was performed in LB (Miller) medium at 30°C or 37°C, respectively, for both liquid and solid agar conditions. As required for *P. putida* propagation*,* 50 mg/L kanamycin, 30 mg/L gentamicin, and/or a combination of 200 mg/L spectinomycin and 300 mg/L streptomycin were used individually for SAGE or combined for mSAGE selection (Supplementary Table S1). 50 mg/L apramycin was used for selection of pGW30 in *P. putida.* For *E. coli*, 50 mg/L kanamycin, 15 mg/L gentamicin, 50 mg/L spectinomycin plus 50 mg/L streptomycin, and 50 mg/L apramycin were used individually for selection. M9 medium (47.8 mM $Na_2HPO_4$, 22 mM $KH_2PO_4$, 8.6 mM NaCl, 1 mM $MgCl_2$, 0.1 mM $CaCl_2$, 18 μM $FeSO_4$, 1x MME trace minerals, pH adjusted to 7 with KOH) with 20 mM $NH_4Cl$ was utilized for shake flask experiments and plate reader cultures. In total, 1000x MME trace mineral stock solution contains per liter, 1 mL concentrated HCl, 0.5 g $Na_4EDTA$, 2 g $FeCl_3$, 0.05 g each $H_3BO_3$, $ZnCl_2$, $CuCl_2{\cdot}2H_2O$, $MnCl_2{\cdot}4H_2O$, $(NH_4)_2MoO_4$, $CoCl_2{\cdot}6H_2O$, and $NiCl_2{\cdot}6H_2O$. The M9 medium was supplemented with 20 mM p-coumarate as a sole carbon source or 20 mM isophthalic acid as the sole carbon sources as required. For soil enrichments, MMM medium was used (Per liter: 7.01 mM $K_2HPO_4{\cdot}3H_2O$, 20 mM 3-(N-morpholino)propanesulfonic acid (MOPS), 4.28 mM NaCl, 9.35 mM $NH_4Cl$, 0.41 mM $MgSO_4{\cdot}7H_2O$, 0.068 mM $CaCl_2{\cdot}2H_2O$, 18 μM $FeSO_4{\cdot}7H_2O$, and 1 mL of 1000x MME trace metal stock solution; pH was adjusted to 7.2 with a 10 M KOH solution). All minimal media were prepared by adding components in the order listed, adjusting pH as described, then filter-sterilizing using a 0.22 micron PES filter.

### mSAGE base strain and plasmid design

Previously reported SAGE tools were modified and expanded to enable multiplexed, simultaneous DNA integration into the chromosome of *P. putida* KT2440. We designed and constructed new plasmids to insert three *attB* sites into the chromosome with ~20 base pair spacer sequences in between to allow for PCR screening to verify integration events. These "landing pads" of *attB* sites were flanked with *rho*-independent terminators [1] to reduce transcriptional read-through into and out of the construct. Recombinase expression 'helper' plasmids (pGW31-33,35-40) were not altered from the original SAGE toolkit [2]. We expanded the functionality of the original integrating *attP* plasmids (pJH204-212) by constructing new plasmids with alternate selectable markers and origins of replication (Supplemental Figure S3). The initial kanamycin resistance gene (*npt*II) was replaced by either a gentamicin resistance gene (*aac(3)-IIa*, pJH213-221) or a spectinomycin resistance gene (*aad*A1, pJH222-230). The ColE1 origin of replication was replaced with either the ColA (pJH213-221) or CloDF13 (pJH222-230) origin of replication to avoid the presence of identical DNA that could allow undesired homologous recombination events within multi-integrated strains. To simplify external vendor DNA synthesis, alternate plasmids (pJH401-427) wherein the pJH204-230 origins of replication were all replaced with the ColE1 origin and a secondary antibiotic resistance gene, *bla*, enabling selection for ampicillin resistance in *E. coli*. Together, this established a suite of plasmids to evaluate and demonstrate mSAGE functionality as a tool for high efficiency DNA integration.

### Plasmid Construction

All PCR reactions were carried out with Phusion® High Fidelity Polymerase (Thermo Scientific) utilizing primers synthesized by Eurofins Genomics. Synthetic DNA fragments and genes were obtained from Integrated DNA Technologies (IDT). For in-house cloning, plasmids were constructed via T4 DNA ligation (New England Biolabs – NEB) or NEBuilder® HiFi DNA Assembly Master Mix (NEB) per the manufacturers standard protocol. All enzymes utilized for plasmid digestion were obtained from NEB. All plasmids were transformed into NEB5-alpha F' IQ competent *Escherichia coli* (NEB) per the manufacturers standard protocol. Transformants were selected on the appropriate LB (Miller) plates containing either 50 mg/L kanamycin, 15 mg/L gentamicin, or 50 mg/L spectinomycin and incubated at 37°C. DNA extraction was carried out using the ZymoPure Miniprep kit from Zymo Research, per the manufacturer's protocols. Plasmids were verified via Sanger sequencing performed by Eurofins Genomics. All plasmids not cloned in-house were synthesized and cloned by Genscript. All plasmids utilized in this work are described in Table S1.

### Strain construction

*P. putida* KT2440 was utilized as the wild-type parent strain for all strains constructed in this work. A base strain with three *att*B landing pads, called AG5577 (WT KT2440 Δ*PP_2876::R4_phiBT1_MR11* Δ*PP_4740::BxBI_RV_phi370* Δ*PP_4217/4218 intergenic::TG1_BL3_A118*), was constructed via the pK18mob-sacB kanamycin selection and sucrose counter-selection system as described previously [3, 4]. In summary, ~3000 ng of plasmid was used to electroporate 50 µL of KT2440 competent cell in a 0.1 cm cuvette at 1.6 kV, 25µF, 200 ohms using a BioRad GenePulser. After a 1 hour recovery in 1 mL of LB at 30˚C at 225 RPM, the recovery was selected overnight on LB plates containing 50 mg/L kanamycin at 30°C. Transformants were single colony purified on LB plates containing 50 mg/L kanamycin at 30°C to ensure untransformed cells were not carried over. Counter-selection was achieved by streaking colonies onto YT+25% sucrose plates (10 g/L yeast extract, 20 g/L tryptone, 250 g/L sucrose and 18 g/L agar) and incubated overnight at 30°C. The resulting colonies were screened via PCR for the excision of plasmid backbone and insertion of the desired poly-*att*B landing pad. PCR-correct colonies were cultured overnight at 30°C, fully verified with 3 additional PCRs and mixed 1:1 with 50% glycerol for freezer storage. The landing pads were inserted sequentially using 1) pGW98, 2) pGW97, and 3) pGW99.

To construct strains via serine recombinase integration, 100 ng of genetic cargo *att*P plasmids were pooled with the appropriate recombinase-expression plasmid in the same quantity. Competent cells of *P. putida* KT2440 strains were generated from stationary phase LB cultures that were incubated overnight at 30°C and shaken at 225 rpm. The competent cells were then processed in similar fashion to previously described methods [4, 5]. Briefly, the cells are washed three times in 10% glycerol at half the original culture volume and then resuspended in 1/50$^{th}$ culture volume of 10% glycerol. To achieve the highest efficiencies and for repeated use, we typically used a 50 mL culture in a shake flask; however, cultures of 5-10 mL in culture tubes can be utilized to generate enough cells for 1-2 transformations. The pooled plasmids were then added to 50 μL of competent cells and electroporated in the same manner as previously described. Due to the high efficiency of integration, 5% or less of the 1000 μL recovery mixture was plated onto LB plates containing the appropriate combinations of 50 mg/L kanamycin, 30 mg/L gentamicin, and 200 mg/L spectinomycin plus 300 mg/L streptomycin and incubated at 30°C overnight. Single colony isolates were screened for correct integration as needed via PCR. PCR-correct colonies were cultured overnight at 30°C in LB medium containing appropriate antibiotics as concentrations previously listed. The strains were then fully verified with 3 additional PCRs using primers oJH0716, oJH0717, and oJH0754-oJH0760, which bind in the SAGE cargo vectors and the *P. putida attB* landing pads to amplify unique fragments from the resulting SAGE. Cultures with correct PCR bands were selected and mixed 1:1 with 50% glycerol for freezer storage. All strains utilized in this work are listed in Supplemental Table S1 and oligonucleotides used for strain verification are listed in Supplemental Table S2.

### Transformation assay for integration of single and multiple plasmids into chromosome via SAGE and mSAGE

A co-transformation assay of a single integrating *attP* plasmid and the appropriate recombinase expression plasmid was performed to evaluate the efficiency and accuracy of DNA integration (Supplemental Figure S1A). In triplicate, pJH204-212 were individually transformed with the corresponding non-replicating recombinase-expression plasmid and plated on selective media. For comparison, a replicating plasmid (pBBR1-MCS2) and a homologous recombination plasmid (pGW97) were transformed to provide relative efficiency data for those methods. Competent cells of AG5577 were prepared from a 50 mL overnight LB culture as described previously. 100 ng each of the genetic cargo plasmid and the recombinase helper plasmid (e.g., pJH204 with pGW31) were combined, electroporated, and recovered in the same manner described above. For plating, 1 μL and 10 μL of recovery mixture was diluted in 99 μL and 90 μL SOB, respectively. The entire dilution was then plated on LB media containing the appropriate antibiotic concentrations and incubated overnight at 30°C. Colonies were counted to determine the number of transformants generated and calculate the cfu/ug DNA. For each individual recombinase, twenty isolates were screened via PCR to confirm accurate integration. In the same manner, co-transformation efficiencies were calculated by pooling 100 ng of each of the three genetic cargo plasmids with 100 ng of each of the appropriate recombinase helper plasmids. Following recovery and overnight selection on LB agar plates with 50 mg/L kanamycin, colonies were counted to quantify transformation efficiency, and PCR screening was completed on twenty-four isolates to confirm accurate integration for all three plasmids. Serine recombinases preferentially integrate DNA between their native *att* sites, but are known to potentially integrate into pseudo-*att* sites [6]. Therefore, integration at the expected *att*B site was evaluated by a PCR screen of twenty-four isolates from each transformation using primers oJH0716, oJH0717, and oJH0754-0760 (corresponding to the anticipated SAGE integration site).

The observed high efficiencies of single recombinase DNA integration suggested that it could be possible to co-integrate multiple plasmids simultaneously. We assessed this by pooling equal mass (100 ng) amounts of three integrating *attP* plasmids (pJH204, pJH216, and pJH228) along with their corresponding recombinase helper plasmids (pGW31, pGW38, and pGW39), totaling 6 plasmids in the pool. This pool was transformed in the same manner as single recombinases and plated on selective media with all required antibiotics. Colonies were counted to quantify the efficiency of simultaneous co-integration. Further, integration accuracy was evaluated via PCR screening of twenty-four isolates using primers oJH0716-0721 and oJH0754-0760.

### Plasmid Backbone Excision

PhiC31 recombinase and other large serine recombinases perform recombination at two specific base pairs near the center of the *att* sites [7]. These two bases must be identical between the *attB* and *attP* sites and coordinately changing these two bases results in orthogonal pairs of *att* sites. We used this concept to create plasmid backbones that can be removed simultaneously without risk of cross-reactivity by changing the second *attB*/*attP* pair to have a GT core and the third to have TC.

A PhiC31 recombinase-expression plasmid (pGW30) was used that contains an origin of replication previously described to show temperature sensitivity in *P. aeruginosa* and *Pseudomonas fluorescens* [8, 9].

This plasmid provides resistance to 50 mg/L apramycin at temperatures ≤ 30°C. To excise plasmid backbones following integration, competent cells of strains were cultured in the same manner described above. 50 ng of pGW30 was electroporated into 50 μL of competent cells and then mixed with 950 μL of SOB, then recovered at room temperature for 2 hours. Following recovery, 50 μL of the mixture was plated on LB media containing 50 mg/L apramycin and incubated overnight at room temperature. Single colony isolates were then cultured in 1 mL LB and incubated at 37°C while shaking for 4 hours. 50 μL of culture was plated onto LB plates containing no antibiotic and incubated overnight at 30°C. Resulting isolates were then screened by PCR using primers oJH754, oJH755, oJH757, oJH758, oJH759, and oJH760 to verified for SAGE plasmid backbone removal and screened for apramycin sensitivity to confirm loss of the temperature-sensitive plasmid.

### Isophthalic acid pathway ortholog identification and plasmid construction

Initial IPA transport and catabolism genes were selected from the previously described pathway in *Comamonas* sp. strain E6 [10]. The gene sequences encoding *iphABCD* were obtained from the published genome (GenBank accession number BBXH01000000). The corresponding protein sequences were utilized to identify additional orthologs using TheSeed (https://www.theseed.org/wiki/Home_of_the_SEED) [11, 12]. From this search, two orthologous pathways were identified in *Burkholderia* sp. CCGE1002 (GenBank accession numbers CP002013, CP002014, CP002015, and CP002016) and *Maritimibacter alkaliphilus* HTCC2654 (ENA accession number GCA_000152805)[13, 14]. Additionally, three organisms isolated from environmental enrichments based on their ability to grow on IPA as a sole carbon and energy source were mined for orthologous IPA catabolic pathways. To isolate these organisms, soil samples were enriched using MMM medium supplemented with 10 mM isophthalate and incubated at 30˚C, 225 RPM. After 1 week, the enrichment was streaked on an MMM agar plate with 10 mM isophthalate and incubated at 30˚C until single colonies appeared. Colonies were restreaked for isolation, then used to inoculate 10 mL of MMM with 10 mM isophthalate. Cell pellets from this culture were harvested by centrifugation at 4000 x g and resuspended in 500 μL of RNAprotect. Genomic DNA extraction and whole-genome resequencing was performed by SNPsaurus. Genome assemblies were annotated by RAST annotations.

Theses isolates were found to possess putative operons with sequence similarity and genomic context similar to *Comamonas* sp. E6. These isolates included organisms with 16S sequences similar to *Comamonas testosteroni, Acidovorax wautersii*, and *Paraburkholderia tuberum*. *P. tuberum* encodes orthologs of *ipaABD*, but no *ipaC* transporter homolog was identified. Instead, a nearby putative MFS transporter was included in the library as a potential IPA transporter. The details of the orthologs can be found in Supplementary Table S3. Using the translated protein sequences from these organisms for *iph*ABCD orthologs, new gene sequences were created to remove common restriction enzyme sites and harmonize codon distributions in *P. putida* using Geneious software tools. Using these sequences, novel plasmids were synthesized into mSAGE based backbones. Orthologs of *iphA* and *iphD* were oriented in an operon, under the control of the $P_{tac}$ promoter and JER01 RBS sequence previously described [3] and were synthesized and cloned into the mSAGE plasmid pJH401 digested with BamHI and XbaI. Orthologs of *iph*B were constructed in the same manner as *iphAD* plasmids except using the mSAGE plasmid pJH413. A slightly weaker promoter variant (pJE111411) was used for the expression of the membrane-associated transporter genes, and these genes were assembled into the mSAGE plasmid pJH425, resulting in plasmids pJH465-481, which were synthesized, constructed, and sequence verified by Genscript before delivery.

### IPA pathway optimization

To build the combinatorial library for IPA metabolism, we combined approximately 500 ng each of pJH465-481 with 5000 ng each of pGW31, pGW38, and pGW39. From this pool, 5 μL of the mixture of plasmids was added to 50 μL of AG5577 competent cells and the electroporation was carried out as described above. Following the recovery period, the entire recovery mixture, less the volume plated for CFU determination, was used to inoculate a 50 mL culture of liquid LB media with 50 mg/L kanamycin, 30 mg/L gentamicin, 200 mg/L spectinomycin, and 300 mg/L streptomycin to create a mixed pool of triply inserted strains. This culture was incubated overnight at 30°C while shaking at 225 rpm. The resulting population was then centrifuged in a 50 mL conical tube at 3000 rpm for 15 minutes to pellet the cells. The entire population was then gently resuspended in 50 mL of M9 minimal media with p-coumaric acid (ThermoFisher #A15167) as the sole carbon source. This culture was then incubated overnight at 30°C while shaking at 225 rpm. Following this incubation, the entire population was pelleted in the same manner and the resulting cell pellet was gently resuspended in 1x M9 salts. From this suspension, 300 μL were transferred to three 50 mL M9 cultures with IPA as the sole carbon source and incubated at 30°C while shaking at 225 rpm. Similarly, 6 μL of the suspension was transferred to a 48 well microtiter plate with 600 μL of M9 medium with IPA as the sole carbon source and incubated at 30°C in a Biotek Epoch 2 plate reader. Initially, all cultures were incubated for approximately 72 hours, then sub-cultured and incubated for 48 hours under the same conditions three more times. From all cultures, samples were collected and a whole DNA extraction was performed using a Zymo Quick-DNA Bacterial miniprep kit (Zymo, D6005). Along with this, samples were collected and mixed with 50% glycerol to be stored at -80°C.

### Ortholog Population Sequencing

To evaluate the composition of the ortholog populations over the entire experimental process, a targeted PCR amplification was performed. Using a common forward primer (oJH966) in equimolar amount to the total molar amount of 17 ortholog specific primers (oJH967-83), a pooled PCR was performed to amplify an approximately 240 bp fragment. A Tm of 56˚C was used for primer annealing, and the PCR reaction was run for 25 cycles. Following amplification, the individual reactions were size selected and isolated from a 2% agarose gel using the ThermoFisher GeneJET gel extraction kit (K0691). The purified amplicons were then prepared for Illumina sequencing using the NEBNext Ultra II DNA library prep kit (NEB #E7645) per the manufacturer's instructions. Briefly, the amplicons were end repaired and ligated to NEBNext adaptors. Following ligation, the amplicons were size selected using magnetic beads. Using barcoded primers, the amplicons were again amplified for each time point and replicate using a unique barcode set. The barcoded amplicons were then sequenced using an Illumina MiSeq instrument.

### Reconstruction of individual pathway combinations

To validate this methodology, ortholog combinations mirroring those we identified through the enrichment process were re-constructed individually. Initially, two best orthologs (e.g., the *A. wautersii* dioxygenase and *Comamonas* sp. E6 dehydrogenase) were integrated into AG5577 to create a strain that could be used to test each transporter. Similar strains were created to test the dioxygenases and dehydrogenases. Each ortholog of the third pathway component (transporter, in the first example) was then integrated to create strains harboring two top components of the optimized pathway paired with each the individual orthologs of the third component not identified by optimization. This allowed for targeted comparison of growth characteristics for the orthologs of optimized genetic combinations against those not enriched to evaluate our selective optimization.
